# Maize kernel composition and morphology influences pericarp retention during nixtamalization

**DOI:** 10.1101/2025.06.29.662213

**Authors:** Michael J. Burns, Sydney P. Berry, Amanda M. Gilbert, Peter J. Hermanson, Candice N. Hirsch

## Abstract

**Background and Objectives:** Pericarp retention during nixtamalization directly influences masa quality, affecting texture, machinability, and nutritional content of staple foods such as tortillas and chips. Despite its industrial relevance, the underlying kernel traits that govern pericarp retention remain poorly characterized. This study aimed to assess an existing staining method to quantify pericarp retention based on visual evaluation, and identify the compositional and morphological characteristics of maize kernels that most strongly predict pericarp retention during nixtamalization.

**Findings:** Stain-based scoring of pericarp retention showed moderate correlation with directly measured pericarp mass, achieving a Pearson’s correlation coefficient of 0.77 in the rapid cook test and 0.56 in the benchtop cook test. Pearson correlation coefficients of 18 compositional and morphological traits range from -0.294 to 0.538. Ground kernel ash content and initial pericarp quantity were the most correlated and predictive variables associated with pericarp retention.

**Conclusions:** Stain-based methods lack the resolution necessary for quantitative pericarp assessment. Initial pericarp quantity and ground kernel ash content are likely key determinants of nixtamalization pericarp retention.

**Significance and Novelty:** This study provides a first assessment on the impacts of morphological and compositional variation for pericarp retention, which can be built upon to develop improved varieties and cooking methods for product optimization.

## 1 INTRODUCTION

Masa is a maize dough created through a high-heat, high-pH cooking process called nixtamalization which loosens the pericarp and softens the endosperm (Santiago-Ramos et al., 2018; Serna-Saldivar et al., 2008). Masa-based products, such as tortillas and tortilla chips are a significant source of calories, calcium, and niacin throughout many parts of the world (Acosta-Estrada et al., 2023; Carpenter, 1983; Serna-Saldivar & Chuck-Hernandez, 2019). Masa quality depends on several factors such as moisture content, texture, nutrition, and 6machinability (Holmes et al., 2019; Miranda et al., 2013). Though the ideal value for each of these traits depends on the final product (i.e. tortilla chips, tortillas, tamales, masa harina), each is affected by the quantity of pericarp that is retained in the masa dough after nixtamalization.

The pericarp is the outermost layer of tissue on a kernel and is the first reactant for the alkaline solution (Iribarren et al., 2002; Martínez et al., 2001). During nixtamalization, the maize pericarp becomes imbibed with water containing Ca(OH)_2_, which begins breaking down the hemicellulose of the pericarp (Santiago-Ramos et al., 2018). The alkaline hydrolysis creates holes in the pericarp that loosen and seperate the it from the kernel, allowing the alkaline solution to permeate the endosperm (Iribarren et al., 2002; Santiago-Ramos et al., 2018). This degradation and removal of the pericarp allows the alkaline solution into the endosperm of the kernel. This softens the grain for future grinding into the masa dough and fortifies the kernels with bioavailable calcium. The alkaline solution also releases starch-bound niacin which fortifies the masa with vitamin B_3_ (Carpenter, 1983). The pericarp is a concentrated source of fibers and gums in the kernel (Martínez-Bustos et al., 2001). If too much pericarp is removed during nixtamalization, the dough will not be cohesive enough and will crumble (Martínez-Bustos et al., 2001; Serna-Saldivar & Chuck-Hernandez, 2019). Conversely, if too much pericarp remains the dough can be overly adhesive causing it to stick to machinery and to sheet improperly (Martínez-Bustos et al., 2001).

Manufacturers can adjust cooking parameters to adjust pericarp retention, but many of these adjustments are made on a post hoc basis. Nixtamalization parameters such as cooking and steeping time, cooking and steeping temperature, pH, and water spray pressure and volume all have an inverse relationship with the quantity of pericarp retained by kernels (Rooney & Serna-Saldivar, 2015). Consistent cooking parameters that minimize input resources would be ideal for manufacturers to limit both reagent input and waste output. If manufacturers had a rapid and accurate method for testing pericarp retention before cooking, or a line of maize that was bred for ideal pericarp retention values according to their nixtamalization parameters, the nixtamalization process could be more efficient with regards to pericarp removal.

Understanding the aspects of a maize kernel that are key determinants of pericarp retention is necessary to make cooking process and germplasm improvements. Base hydrolysis is the accepted method by which pericarp is degraded in nixtamalization (Santiago-Ramos et al., 2018), but it is unclear to what extent the chemical composition of the pericarp, initial pericarp quantity, or kernel shape impact pericarp retention. The composition of the pericarp could impact its retention given that some hemicellulose binders or stabilizers could be more recalcitrant to base hydrolysis than others (Serrano-Gamboa et al., 2019). Initial pericarp quantity could also govern pericarp retention by providing more substrate that needs to be degraded before further imbibition can occur. Even kernel shape can theoretically impact pericarp retention by changing the surface area to volume ratio, the thickness of pericarp at any given point in the kernel, and the packing density of kernels in a cooking vessel.

Understanding the contribution of chemical and morphological characteristics requires robust methods of measurement. Compositional assessments can be performed using near-infrared (NIR) spectroscopy, which is a high-throughput method of determining grain composition (Renk et al., 2021; Varela et al., 2022). Likewise, image analysis methods have been developed to assess kernel shape characteristics in a high-throughput manner (Miller et al., 2017; Wang et al., 2025). Determining initial pericarp quantity requires meticulous peeling from kernels and is a low-throughput evaluation method (Cope et al., 2006). With regards to pericarp removal, a high-throughput evaluation method for the ease of pericarp removal was previously developed that boils kernels in a 0.5% Ca(OH)_2_ solution for 20 minutes before staining them with a methylene blue and eosin y solution to create contrast between the pericarp and the aleurone (Serna-Saldivar et al., 1991). The kernels are rated on a stain coverage scale of 1 to 5 where each integer increase corresponds to a 20 percent increase in stain coverage. This method, termed here the rapid cook test, has been used in several studies to assess food-grade maize samples for their ease of pericarp removal (Bhatnagar et al., 2004; Pozo-Insfran et al., 2007; Serna-Saldivar et al., 2008). However, the method, which was developed to stain and quantify the pericarp and endosperm fractions of milled sorghum (Scheuring & Rooney, 1979), does not provide a direct quantification of pericarp retention.

Here, we seek to understand how different aspects of the grain composition and morphology impact the pericarp retention of maize kernels. To this end, we optimized the staining procedure in the previously described rapid cook test to create stronger contrast between pericarp and aleurone and evaluated this method for its use in assessing pericarp retention. The relationship between compositional and morphological traits with pericarp retention was investigated in a diverse set of maize hybrids, and the most important traits based on correlation and model contribution were validated in an independent set of genotypes. This study provides foundational information on the contribution of maize kernel composition and morphology that are associated with pericarp retention. This information provides a biological basis for future study to optimize the removal of pericarp during the nixtamalization process.

## 2 MATERIALS AND METHODS

### 2.1 Plant materials

A panel of 819 diverse hybrids were created from crossing B73, Mo17, and LH244 as the pollen donor to ∼280 inbred lines of the Wisconsin Diversity Panel (Burns et al., 2025; Hansey et al., 2011; Supplemental Table S1). The hybrids generated from these crosses were planted at the Agricultural Experiment Station in St. Paul, MN in Summer 2022 and 2023. The experiment included two replicates each year in a randomized complete block design where early and late flowering genotypes were blocked within replicates. Grain was collected from each plot and dried in a 63°C forced air oven for seven days.

Grain harvested from two potential commercial hybrids (coded as P1306 and P1306W) by Pioneer Hi-Bred International, now Corteva Agriscience, were used to optimize the staining protocol described below.

### 2.2 Spectral and compositional data collection and analysis

Spectral and compositional profiles for each experimental entry was obtained through two near-infrared (NIR) spectroscopic methods. The first method used a Perten DA7250 to obtain 141 wavebands from 950nm to 1650nm at 5nm intervals from ground grain samples as previously described (Burns et al., 2025). The second method used a FOSS Infratec Nova to scan ∼110g of whole kernel samples to obtain 1400 wavebands from 400nm to 1099.5nm at 0.5nm intervals. The FOSS Infratec Nova sample transport module (STM) was utilized to reduce the required grain quantity for each sample. The number of subsample scans was set to five and the sample path length was set to 29mm. The spectra from the FOSS Infratec Nova was downloaded as a .nir file which was converted to a .csv format using the Spectragryph software v1.2.16 (Menges, 2022).

Compositional data was predicted from both the Perten and FOSS NIRs using previously developed equations. The Perten global compositional profiles included protein, starch, fiber, fat, ash, and moisture on an as-is basis and were moisture corrected as previously described (Burns et al., 2025). The compositional prediction equations developed by Perten are based on proprietary models with over two-thousand training samples. The FOSS local compositional profiles (Parvej et al., 2020) included protein, starch, oil, and moisture. The FOSS compositional data was provided on a moisture corrected basis and was not corrected post hoc.

The spectra from the FOSS Infratec Nova was used to create an initial dataset of 60 spectrally diverse samples that were used to assess the rapid cook test methodology against the benchtop cook test methodology and to assess the relationship between compositional and morphological traits with pericarp retention. To identify the most spectrally diverse samples, the spectral data was downsampled to every fifth waveband from 400nm to 1095nm, and normalized through standard normal variate decomposition in the prostpectr v0.2.4 package (Stevens & Ramirez-Lopez, 2025). Honigs regression (Honigs et al., 1985) was used on the normalized spectra to find 12 spectrally diverse samples from each pollen parent (N=3) and year (N=2) combination that did not have overlapping egg parents (Supplemental Table S1).

### 2.3 Nixtamalization protocols

The 60 spectrally diverse samples were cooked in replicate following a previously described rapid cook test method (Serna-Saldivar et al., 1991) and a previously described benchtop cook test that more closely represents the nixtamalization process that occurs at manufacturing scales (Burns et al., 2021; Yglesias et al., 2005).

The rapid cook test was scaled down from the original method as follows. A 4L beaker with 3500ml of 0.6% Ca(OH)_2_ was heated on a hotplate set to 400°C. Forty kernels per sample were placed in 10.16cm by 11.98cm organza bags. Thirty bags of kernels were strung over a plastic rod to make manual agitation more effective. The samples were placed in the beaker of boiling Ca(OH)_2_ solution with the plastic rod placed across the opening of the beaker and covered with tinfoil to reduce evaporation and subsequent concentration of the cooking liquor. The samples were cooked for 20 minutes and were agitated every four minutes by lifting and lowering the samples 5 times followed by 10 seconds of stirring with a glass rod. Following the cook, the plastic rod holding the samples was lifted and samples were placed in a 4L beaker containing cold tap water to halt further cooking. The samples were then placed under running tap water for 3 seconds each to wash away loose pericarp. A subsample of 20 kernels were placed in a metal strainer which was submerged in an optimized stain solution (described below in Section 2.5) for 15 seconds followed by three 10 second washes in methanol. Stained kernels were then rated on a scale of one to five as previously described (Serna-Saldivar et al., 1991), where each integer increase corresponds to a 20% increase in the surface area of kernels covered by pericarp(i.e. 1=0-20%, 2=20-40%, 3=40-60%, 4=60-80%, 5=80-100%). A second subset of 20 kernels were split into two ten-kernel replicates which had the retained pericarp manually removed to quantify pericarp retention (described below in section 2.4).

As with the rapid cook test, following the benchtop cook test protocol (Burns et al., 2021), two subsets of 10 kernels underwent manual pericarp removal for quantifying pericarp retention and 20 kernels underwent staining and scoring on the same 1-5 scale.

### 2.4 Quantification of pericarp retention

From a given sample, two subsets of 10 randomly chosen kernels had the retained pericarp manually removed to quantify pericarp retention. Pericarp was removed from each kernel using forceps, with care not to remove tip cap, aleurone, germ, or endosperm from the kernel. The pericarp and cleaned kernels were placed in separate tin foil containers, covered, and dried in a forced air oven set to 103°C. The desiccated pericarp samples were weighed after two days in the oven and the desiccated kernel samples were weighed after seven days. The mass of pericarp was determined by subtracting the mass of the weighing dish from the mass of the weighing dish with pericarp. The kernel mass was determined by weighing the desiccated kernels directly. The dry mass of pericarp was divided by the dry mass of kernels to normalize for kernel size as larger kernels would inherently have more pericarp if pericarp thickness was held constant.

### 2.5 Stain optimization

The stain previously proposed for contrasting pericarp from endosperm or aleurone was the May-Grunwald stain (Scheuring & Rooney, 1979; Serna-Saldivar et al., 1991) which is 0.5% w/v methylene blue and eosin y in methanol with three washes in methanol after staining. The methylene blue stains the pericarp blue while the eosin y stains the endosperm dark pink. To further optimize the staining protocol, the stain solvent polarity, the components of the stain, the concentration of the components, and the wash solvent polarity were tested on a yellow hybrid (P1306) and a white hybrid (P1306W). The intensity of color imparted on the pericarp, and ease of stain removal from the aleurone (contrast) were used to determine the ideal staining protocol. The stain solvent polarity was tested by staining nixtamalized kernels with methylene blue and eosin y in methanol or deionized water and washed with methanol. The methylene blue and eosin y were tested separately and combined at 0.5% w/v concentrations in deionized water and washed with methanol. The minimum necessary concentration of methylene blue was tested at concentrations of 0.25%, 0.5%, and 0.75% with methanol washes. Finally, the wash solvent polarity was tested by comparing methanol washes to deionized water washes to determine which solvent would minimize overstaining. The ideal staining protocol was 0.5% w/v methylene blue in deionized water with three methanol washes, and was used for all staining.

### 2.6 Measuring kernel morphology traits

Image analysis was used to capture multiple kernel morphological traits. Images were collected from approximately 80 kernels of the grain used in the benchtop cook test before nixtamalization occurred by adapting a previously described image capture protocol (Miller et al., 2017). Briefly, an Epson V800 flatbed scanner was used to collect images through the VueScan software v9.7.65 (Hamrick Software, 2021) with the resolution set to 1200dpi and no color correction utilized. Files were saved in TIFF format due to the high resolution and compression fidelity. The backdrop of the kernel images was uniform black paper. Images were processed in Python v3.9.6 (Van Rossum & Drake, 2009) using scikit-image v0.18.1 (van der Walt et al., 2014). Images were converted from the red-green-blue color space to grayscale with values from 0 to 1 and masked using a threshold of 0.3. Pixels above this threshold were considered kernels, and pixels below the threshold were considered background. Masked objects smaller than 5,000 pixels were removed from the image and holes smaller than 5,000 pixels were filled in through dilation. Each kernel in the masked image was given a unique label, and the shape parameters were determined using the regionprops_table function. The shape parameters extracted were area, perimeter, major axis length (kernel length), minor axis length (kernel width), eccentricity (circularity), and extent (rectangularity). These measurements were then averaged across the ∼80 imaged kernels from the sample.

Additional kernel features were manually collected, including: kernel mass, kernel volume, kernel density, and initial pericarp quantity. Kernel mass was determined with a four-digit scale by weighing 15 random kernels from a given genotype. These kernels were submerged in water in a graduated cylinder, where the water displacement was recorded as volume. The kernel density was calculated from the kernel mass and volume. The kernels were then imbibed in water for 16 to 24 hours and five kernels were manually peeled with forceps to determine initial pericarp quantity. The peeled pericarp and remaining kernels were placed in separate weigh boats and dried in a 103°C oven. Dry pericarp mass was measured after two days in the oven and dry kernel mass was measured after seven days in the oven. The mass of dried pericarp was normalized for kernel size by dividing by the mass of dried kernels to obtain the initial pericarp quantity.

### 2.7 Comparison of the cook test methods and the stain protocol for measuring pericarp retention

The mass of pericarp retained from the rapid cook test, mass of pericarp retained in the benchtop cook test, and the stain ratings from each method were compared. Pericarp retention for both the rapid cook and benchtop cook were normalized through a logarithmic transformation due to the right-skewed distribution of values and the increasing variance that occurred with larger phenotypic values. The 1-5 stain rating scale was not normalized so as to not alter the relationship between the score and percent coverage. Pearson’s correlation coefficient was calculated between the observed pericarp retention in the rapid cook test and the associated stain rating, the observed pericarp retention in the benchtop cook test and the associated stain rating, the pericarp retention of the rapid cook test and the benchtop cook test, and the stain ratings across cook test methods.

### 2.8 Relationship of kernel composition and morphology with pericarp retention

Correlation analysis was first used to test the relationship between pericarp retention and compositional and morphological traits using the 60 spectrally diverse samples. The benchtop cook test is the closest estimate of pericarp retention and was used for this analysis. Compositional and morphological data were collected through NIR spectroscopy, image analysis, and manual kernel measurements as described above. Correlation was calculated using Pearson’s correlation coefficient and significance was corrected for using Bonferroni correction.

To further assess the predictive relationship these traits have with benchtop cook test pericarp retention, a lasso and elastic net regularized generalized linear regression (glmnet) model was created with the caret package v6.0-86 (Kuhn, 2008) in R v4.0.3 (R Core Team, 2022). The glmnet model was fit to predict benchtop cook test pericarp retention from all compositional and morphological traits in a five-fold cross validation training scheme. The feature importance was extracted using the varImp function in caret v6.0-86 (Kuhn, 2008) with scaling turned off. Due to the variance associated with training a model under cross validation, the model training and feature importance extraction were performed 100 times. The feature importance for each feature was determined by averaging its importance across all 100 iterations.

The most promising kernel attributes (initial pericarp quantity, ground kernel ash content, kernel circularity, ground kernel fiber content, and whole kernel protein content) were selected for validation as they had both significant correlations with benchtop cook test pericarp retention as well as showed a high degree of importance to the glmnet prediction model.

### 2.9 Validating the relationship between grain compositional traits and pericarp retention

An independent set of hybrids grown in 2022 and not included in the initial set of 60 spectrally diverse hybrids were selected for validating the relationships of ground kernel ash content, whole kernel protein content, and ground kernel fiber content with pericarp retention. The samples for compositional validation included ten samples from each pollen parent (B73 and Mo17) that were in the upper tail and ten that were in the lower tail of each respective trait distribution and did not have overlapping egg parents across tails, pollen parents, or the initial set of 60 hybrids. Grain from these samples was cooked in replicate using the benchtop cook protocol described above and pericarp retention was measured by manual peeling and weighing the dried pericarp after cooking. A two-sample t-test was performed for benchtop cook test pericarp retention between the high and low groups for each compositional trait.

### 2.10 Validating the relationship between kernel morphological traits and pericarp retention

Measuring initial pericarp quantity and kernel shape parameters is prohibitively time consuming to measure on the full population to select extreme samples, as was done with the compositional trait validation, and as such a different approach was required. To validate the relationship of pericarp retention with initial pericarp quantity and kernel circularity, 200 spectrally diverse samples from the Wisconsin Diversity Panel hybrids were identified through Honigs regression (Honigs et al., 1985). Spectral data from the FOSS Infratec Nova was used for selecting these 200 samples. The set included 40 hybrids from each pollen-parent-year combination that did not have overlapping egg parents with each other or the initial set of 60 hybrids. Initial pericarp quantity and kernel morphology was measured on these 200 samples as described above. From the 200 samples, 40 samples that evenly spread across the distribution of initial pericarp quantity and kernel circularity while maintaining equal representation from each pollen-parent-year combination were selected for benchtop cook test pericarp retention analysis. The correlation of pericarp retention with initial pericarp quantity and kernel circularity was assessed through Pearson’s correlation coefficient to validate the relationship.

### 2.11 Code Availability

The code for this manuscript is available at https://github.com/HirschLabUMN/Pericarp_Retention. The data that supports the findings of this study are available in the supplementary material of this article.

## 3 RESULTS AND DISCUSSION

### 3.1 Stain optimization increased contrast for improved rating of pericarp retained after nixtamalization

Previous literature indicated that the May-Grunwald stain is particularly effective at differentiating bran from endosperm tissues due to the presence of both methylene blue for staining pericarp and eosin y for staining endosperm (Scheuring & Rooney, 1979). Given that nixtamalization does not impact the aleurone (Santiago-Ramos et al., 2018), it is unlikely that the eosin y is beneficial to the stain solution. In addition, kernels are nixtamalized in an aqueous solution and stained in a methanol solution. The difference in polarity between the water in the pericarp and methanol in the stain could impact stain deposition and removal by reducing miscibility (Kunerth, 1922).

To test the utility of the staining protocol and further optimize it, the impact of stain components, solvent polarity, and wash polarity on the contrast created during the previously developed staining protocol (Serna-Saldivar et al., 1991) were assessed with a yellow maize hybrid (P1306) and a white maize hybrid (P1306W). Nixtamalized kernels (Figure 1A) were stained with the proposed May-Grunwald stain with both methylene blue and eosin y at 0.5% w/v concentrations in methanol (Figure 1B) and in water (Figure 1C). The aqueous solvent increased the stain intensity considerably, demonstrating that the water-based stain more effectively deposits stain into the pericarp of the kernels. To test the necessity of both stain components, separate 0.5% w/v methylene blue in water (Figure 1D) and 0.5% w/v eosin y in water (Figure 1E) solutions were tested. The contrast generated by the methylene blue solution (Figure 1D) was significantly more pronounced than the contrast generated by the eosin y solution (Figure 1E), particularly on the yellow hybrid, indicating that methylene blue is sufficient for generating contrast between pericarp and aleurone. To determine the necessary concentrations of methylene blue, 0.25% w/v (Figure 1F) and 0.75% w/v (Figure 1G) stains were tested. The 0.25% w/v solution generated notably less contrast than either the 0.5% or 0.75% solutions, while the 0.75% solution generated deep staining that turns almost black, but also leaves stain behind on kernels after washing, indicating that 0.5% is an ideal concentration for staining pericarp. The washing solution was also assessed given the notable difference in stain deposition between methanol and water solvents. Water was tested as an alternative to methanol in the washing of stained kernels (Figure 1H) and showed less stain removal from the surface of nixtamalized kernels and left notable over-staining on the kernels (Figure 1H). For these reasons, 0.5% w/v methylene blue in water with three methanol washes was determined to be the optimal stain for rating pericarp retained after nixtamalization.

**Figure 1.**
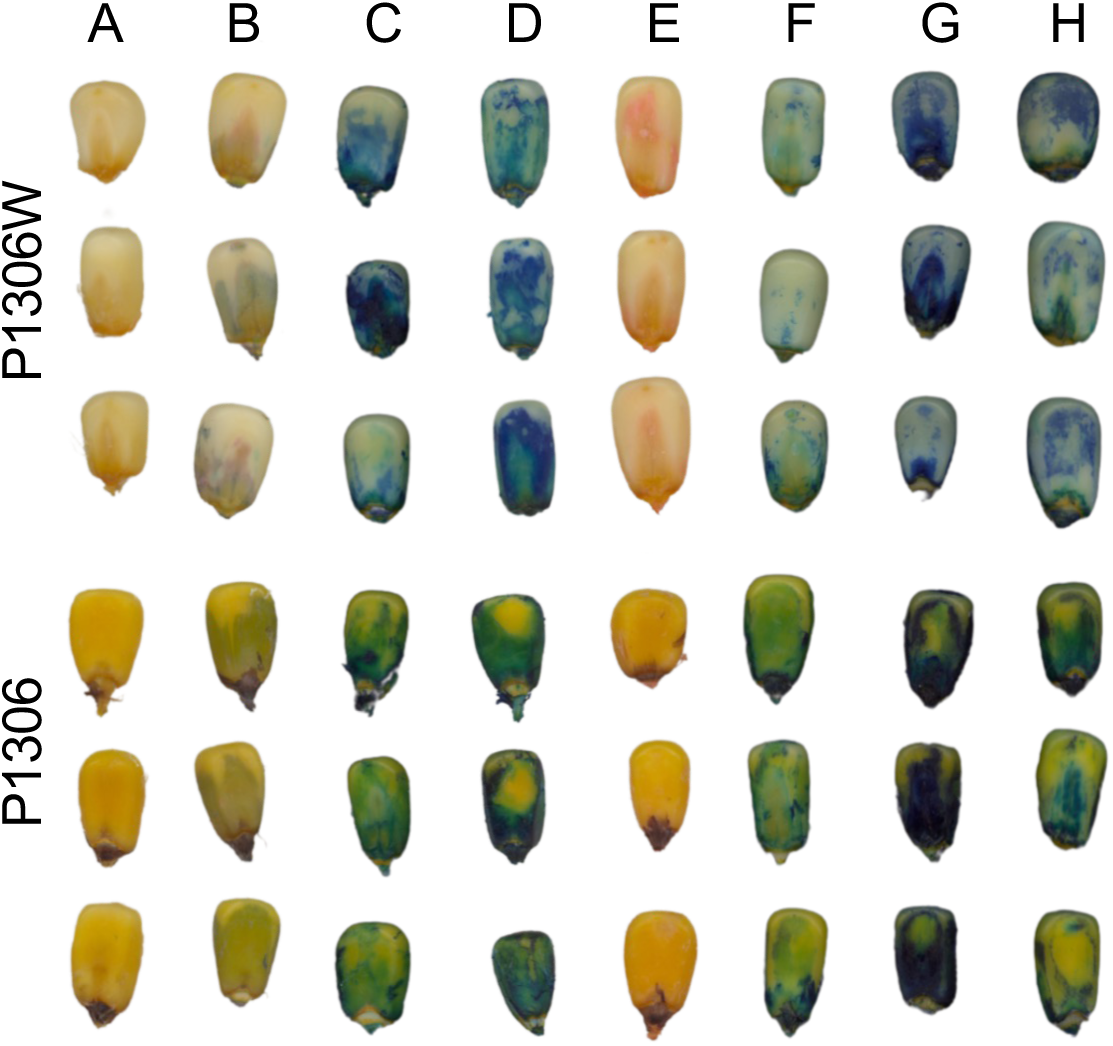
Kernels undergoing stain optimization. (A) Nixtamalized kernels that have not been stained. (B) Kernels stained using 0.5% w/v May-Grunwald stain in methanol with three methanol washes. (C) Kernels stained with 0.5% w/v May-Grunwald stain in deionized water with three methanol washes. (D) Kernels stained with 0.5% w/v methylene blue in deionized water with three methanol washes. (E) Kernels stained with 0.5% w/v eosin y in deionized water with three methanol washes. (F) Kernels stained with 0.25% w/v methylene blue in deionized water with three methanol washes. (G) Kernels stained with 0.75% w/v methylene blue in deionized water with three methanol washes. (H) Kernels stained using 0.5% w/v methylene blue in deionized water with three washes in deionized water.

### 3.2 Spectrally diverse samples are both compositionally and morphologically diverse

While it has been shown that the previously developed rapid cook test and staining method is able to accurately distinguish bran from endosperm tissues (Scheuring & Rooney, 1979; Serna-Saldivar et al., 1991), the relationship between benchtop cook test pericarp retention and stain rating has not been documented. To assess the correlation of benchtop cook test pericarp retention with rapid cook test pericarp retention and stain rating, a representative subset of sixty spectrally diverse samples were chosen from the Wisconsin Diversity panel hybrids (Burns et al., 2025); Supplemental Table S1) based on whole kernel NIR spectra. NIR spectra is a product of both kernel composition (Renk et al., 2021; Varela et al., 2022) as well as the morphology of the kernels (Paponov et al., 2020), thus providing a range of compositional and morphological diversity in the test set. Indeed, the spectrally diverse samples (Supplemental Figure S1) demonstrated diversity in nixtamalization pericarp retention and stain rating, compositional profiles, and kernel morphology (Supplemental Figure S2; Supplemental Table S2). Pericarp retention was right skewed in both cook tests and ranged from 0.65 to 10.94mg of pericarp per gram of kernel in the benchtop cook test and 7.94 to 40.58mg of pericarp per gram of kernel in the rapid cook test. The stain rating was right skewed in the benchtop cook test ranging from one to three, and left skewed in the rapid cook test ranging from two to five. Composition traits were normally distributed in these 60 spectrally diverse samples with protein ranging from 6.12 to 15.64% (ground kernel=6.11 to 15.64, whole kernel=7.23 to 12.4), starch ranging from 62.4 to 80.3% (ground kernel=73.78 to 80.31, whole kernel=62.4 to 66.2), fat ranging from 3.09 to 5.88% (ground kernel=3.33 to 5.88, whole kernel=3.09 to 4.5), fiber ranging from 1.69 to 2.56%, and ash ranging from 1.21 to 1.71% (Supplemental Figure S2). Morphological traits were also normally distributed with kernel mass ranging from 184 to 392mg, kernel volume ranging from 0.15 to 0.32mL, kernel density ranging from 1.08 to 1.35g/L, and initial pericarp quantity ranging from 37.5 to 68.4mg of pericarp per gram of kernel (Supplemental Figure S2).

### 3.3 Rapid and benchtop cook tests are moderately correlated with stain rating

The initial 60 samples were processed in both the rapid cook test (Serna-Saldivar et al., 1991; Supplemental Table S3) and benchtop cook test (Burns et al., 2021; Supplemental Table S4). Subsets of grain from each cook method were processed for manual pericarp retention measurement and stain rating to determine the relationship between the two cooking and two stain-based quantification methods.

Pericarp retention is a difficult trait to measure in a high-throughput and consistent manner as the ground-truth is measured through weighing the mass of pericarp that is peeled off of nixtamalized kernels which can be impacted by both the cooking parameters and the individual performing the removal. To understand the ceiling of correlation that can be expected between cooking methods and quantification methods, the technical variation in this method was first evaluated. The correlation between biological replicates was found to be 0.67 in the benchtop cook test (Supplemental Figure S4A) and 0.75 in the rapid cook test (Supplemental Figure S4B). After further investigation it was found that the person peeling kernels had a large impact on the correlation, particularly in the benchtop cook test. Individual peelers had a correlation above 0.8 between biological replicates when they peeled both replicates. However, when replicates were peeled by different individuals, the correlation was only 0.54, which sets a ceiling on expected correlations with the benchtop cook pericarp retention (Supplemental Figure S5).

The manual and stain-based quantification methods within the rapid cook test were compared next. The Pearson’s correlation coefficient between log transformed rapid cook test pericarp retention and rapid cook test stain rating was 0.77. However, there was a lack of discernment in pericarp retention when the kernel was more than 60% covered (Figure 2A). Next, the same relationship between the manual and stain-based quantification methods was evaluated for the benchtop cook test, which more closely mimics the process used in manufacturing. The Pearson’s correlation coefficient between log transformed benchtop cook test pericarp retention and benchtop cook test stain rating was substantially lower at 0.56 and similarly showed a large overlap between rating groups (Figure 2B). The benchtop cook test, however, showed a lack of discernment at low scoring groups, and had a more limited overall range of observed stained ratings. The lack of dynamic range in stain rating does not fully explain the observed overlap between rating groups. The 1-5 stain rating system is primarily concerned with the proportion of kernel that is covered by pericarp and thus it considers pericarp retention a two-dimensional problem of length and width. In reality, pericarp is a three-dimensional structure and accounting for the thickness is necessary.

**Figure 2.**
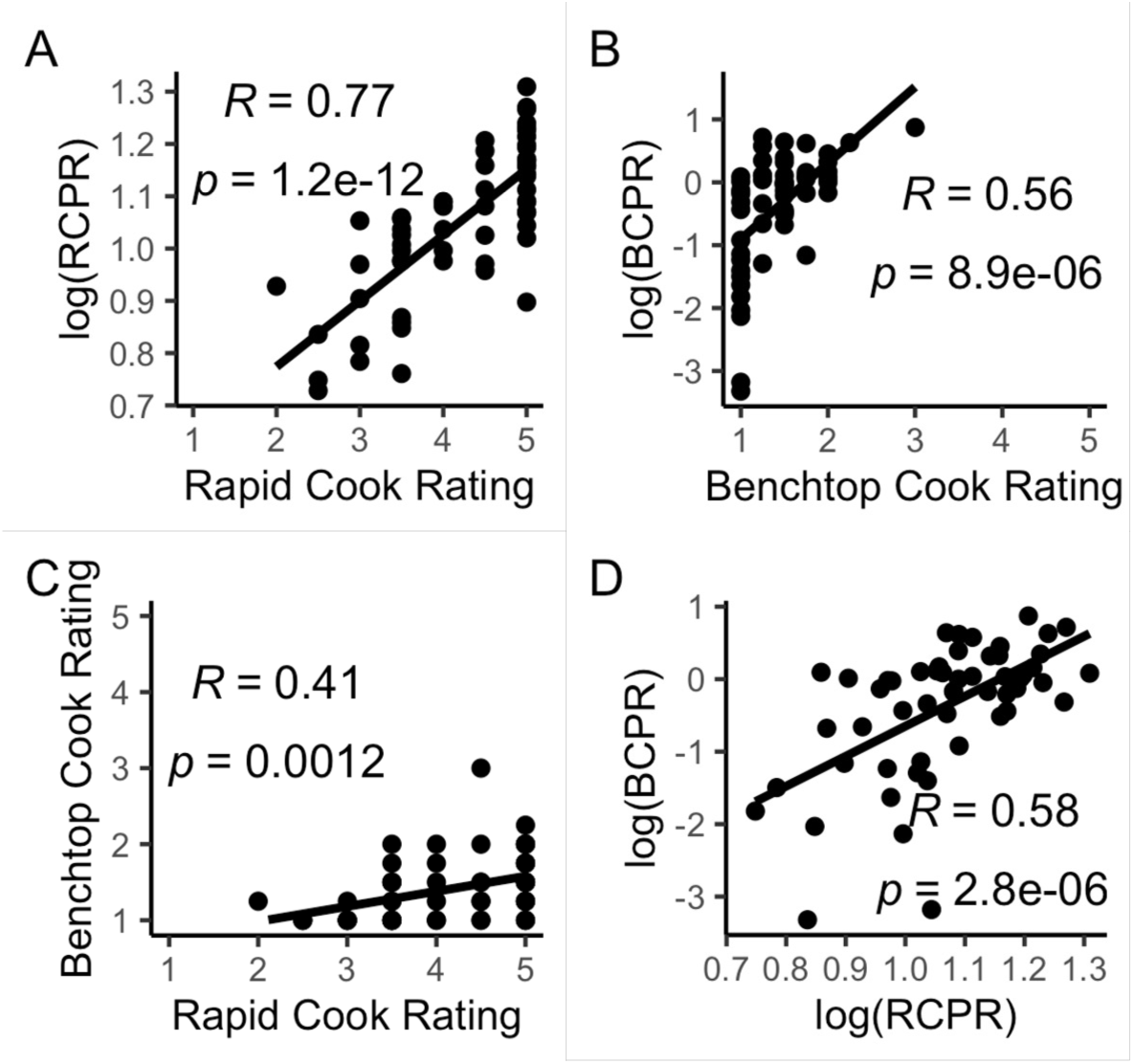
Comparison of cook test methods and stain ratings for spectrally diverse hybrids. (A) Correlation of log transformed rapid cook pericarp retention (RCPR; mg/g) and the rapid cook test stain rating. (B) Correlation of log transformed benchtop cook pericarp retention (BCPR; mg/g) and the benchtop cook test stain rating. (C) Correlation of the stain ratings in the benchtop cook test and rapid cook test methods. (D) Correlation between log transformed benchtop cook pericarp retention (BCPR; mg/g) and log transformed rapid cook pericarp retention (RCPR; mg/g).

Comparisons across cook test methods were also made to determine how representative the rapid cook test is of the benchtop cook test, which is the ideal metric as it most closely mimics the processing that occurs in a manufacturing system (Burns et al., 2021; Yglesias et al., 2005). The Pearson’s correlation coefficient between stain ratings of the rapid cook test and benchtop cook test was 0.41. Most of the rapid cook test stain rated groups overlapped in their benchtop cook test stain rating, again indicating a lack of discernment between rating groups (Figure 2C). The correlation of pericarp retention between the benchtop cook test and the rapid cook test was more significant with a Pearson’s correlation coefficient of 0.58 (Figure 2D). While a moderate correlation between these cooking protocols does exist, care should be taken when using the rapid cook test as a proxy for the benchtop cook test or a manufacturing system.

### 3.4 Compositional and morphological traits associate with benchtop pericarp retention

There are many possible compositional and morphological factors that can contribute to pericarp retention during nixtamalization. To test for potential relationships, 18 different compositional and morphological traits were correlated with benchtop cook test pericarp retention (Figure 3A; Supplemental Table S5). Benchtop pericarp retention had significant correlations with initial pericarp quantity (R=0.538), ground kernel ash content (R=0.479), ground kernel fiber content (R=0.414), whole kernel protein content (R=0.374), kernel circularity (R=-0.320), whole kernel starch content (R=-0.282), ground kernel protein content (R=0.296), ground kernel starch content (R=-0.294) and kernel length (R=-0.255; Figure 3A; Supplemental Table S5). Of these, initial pericarp quantity and ground kernel ash content remained significant after Bonferroni correction (Supplemental Table S5). It is notable that kernel density did not correlate with pericarp retention even though it is one of the proxies used for cooking quality among food-grade maize breeders (Holmes et al., 2019; Rooney & Serna-Saldivar, 2015), demonstrating the importance of measuring traits as directly as possible. The compositional predictions from the Perten NIR spectrometer on ground kernel samples and predictions from the FOSS NIR spectrometer on whole kernel samples were significantly correlated for protein content (R=0.889) and starch content (R=0.592), indicating that the two spectrometers provide similar information, but as they are not perfectly correlated, each can still add unique information (Supplemental Figure S3). While both machines report absorbance, the Perten NIR directly measures reflectance and the FOSS NIR measures transmittance, which are then converted to absorbance before being reported to the user due to the lack of linear relationship between reflectance or transmittance with concentration (Rinnan et al., 2009). The difference between spectrometry methods provide complimentary information about scanned grain samples. For instance, the Perten NIR represents the average composition of the entire grain sample as the grain is ground before scanning and spans the second overtone region of the NIR spectrum (950 to 1650nm), which primarily includes aliphatic, nitrogenous, and hydroxyl groups (Salazar, 2024) that are primarily associated with carbohydrates, proteins, and lipids. Conversely, the FOSS NIR represents external composition as whole kernels were scanned and NIR spectra does not have enough penetrance to measure the entire kernel (Fu et al., 2021). The FOSS NIR spectra spans nearly the entire visible light spectrum into the third overtone region of the NIR spectrum (400 to 1099.5nm; Bock et al., 2020). This portion of the NIR spectrum is largely associated with aliphatic and aromatic functional groups which are primarily associated with proteins and carbohydrates while the visible spectrum provides information about the elemental composition of samples (Nejdl et al., 2022).

**Figure 3.**
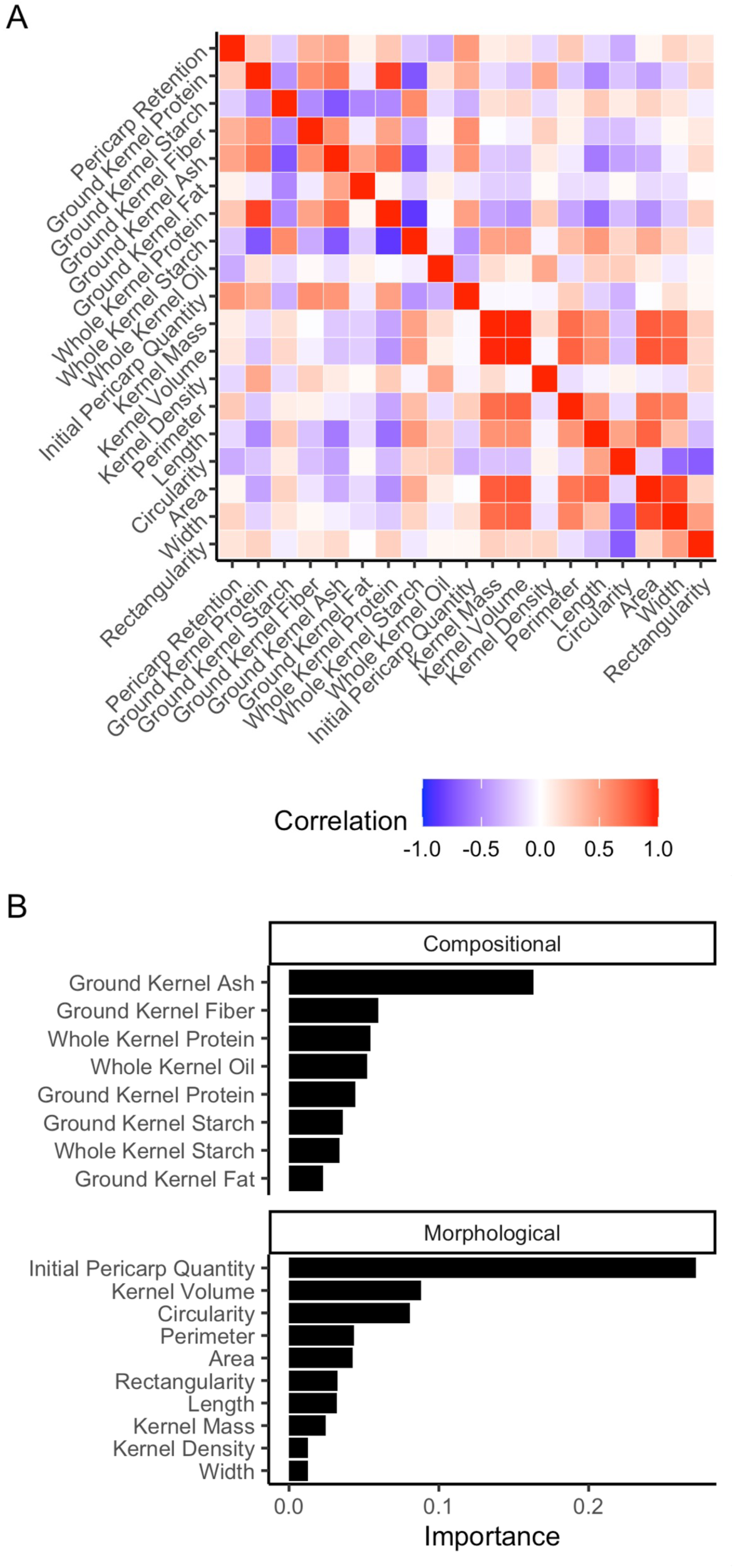
Kernel feature relationships with pericarp retention. (A) Heatmap of Pearson’s correlation coefficients between all traits collected on the 60 spectrally diverse hybrids. (B) Importance of each kernel feature to the lasso and elastic net regularized generalized linear model.

Correlation analysis provides insight into the relationship between two variables in a pairwise process. It does not, however, provide information into how useful many interrelated traits are to predict a dependent variable and cannot account for the intercorrelations of independent variables. Feature selection provides a method to assess the predictive nature of independent variables. It is less likely to provide artifactual results since it reduces correlated traits to single variables when the variables do not provide meaningful additional information to selection models. Thus, to further assess the relationship between benchtop cook test pericarp retention and the compositional or morphological traits, a glmnet model was developed. This model uses L1 regularization to reduce the coefficients, and therefore the relationship between the dependent variable and the independent variables (Heslot et al., 2012). This is a common method of feature selection (Engebretsen & Bohlin, 2019) and can be used to determine which features are most predictive of benchtop cook test pericarp retention. Using this approach, initial pericarp quantity and ground kernel ash content were notably more important than any other feature in predicting benchtop cook test pericarp retention (Figure 3B). These two kernel features were also identified from the correlation analysis (Supplemental Figure S6). Initial pericarp quantity is a measure of the pericarp present on the raw kernel and represents the amount of substrate the alkaline solution needs to react with to fully de-bran a kernel. Ash content is a measure of the mineral content of a sample after incineration and could be related to the chemical recalcitrance of a sample’s pericarp, or could be a proxy of a compositional trait that chemically associates with mineral ions such as certain proteins or functional groups.

In addition to ground kernel ash content and initial pericarp quantity, there was notable glmnet model importance from kernel volume and kernel circularity (Figure 3B). While kernel volume was not significantly correlated with pericarp retention (R=0.014; Supplemental Table S5), kernel circularity was one of the most correlated traits with pericarp retention (R=-0.320) and had just slightly too low of a correlation to reach the threshold set by Bonferroni correction, which is known to be an overly conservative multiple test correction method (Kaler et al., 2019). It is likely that grain circularity impacts the surface area exposed to alkaline solution by decreasing the surface overlap that could occur by more rectangular kernels. In addition, more circular kernels tend to occur when nearby kernels are aborted, meaning that kernels can grow larger than they otherwise would. This increased endosperm size could cause the pericarp to be thinner than it would otherwise be on a smaller kernel by spreading it across a larger surface.

There was a notable decline in glmnet model importance and correlation to pericarp retention from ground kernel ash content to other compositional traits. Many of these traits are intercorrelated (Figure 3A and Supplemental Figure S3), which could impede the ability of the prediction model to attribute feature importance across all of these features. Still, grain protein (ground and whole kernel) and ground kernel fiber content were consistently found among the most correlated to pericarp retention and important features in the glmnet model (Figure 3). Whole and ground kernel protein content are highly correlated (R=0.889), though whole kernel protein was more related to pericarp retention in both correlation and model importance. The increased importance of protein content in the whole kernel spectroscopy could be indicative of the role of binding proteins in the cell wall material of pericarp cells. It could also be indicative of the importance that externally located proteins play in pericarp retention as opposed to internal proteins. Fiber content constitutes both cellulosic and hemicellulosic material, and could be partly measuring the quantity of both materials in a grain sample. The correlation and importance of ground fiber content could be reduced by the fact that kernels and ground before NIR scanning, which allows fibers from inside the kernel to impact the total fiber content even though the internal fibers do not participate in pericarp removal. Protein content could be indicating the role of binding proteins in the cell wall material of pericarp cells. Given that whole kernel protein content was consistently more related to pericarp retention than ground kernel protein content could also be indicative of the importance that externally located proteins play in pericarp retention as opposed to internal proteins.

### 3.5 Samples segregating for compositional traits also segregate for pericarp retention

Based on the correlation analysis and model importance, ground kernel ash content, whole kernel protein content, and ground kernel fiber content were selected from the compositional traits to validate the potential relationships with pericarp retention. To this end, independent sample sets were selected from the tails of each trait distribution (Supplemental Figure S7) and cooked according to the benchtop cook test protocol to measure pericarp retention (Supplemental Table S6 and S7). Due to the discrepancy across peelers (Supplemental Figure S5), the individual peeling pericarp for trait validation was constant throughout the validation of a given trait.

Samples with high ground kernel ash content had an average of 2.38mg of pericarp per gram of kernel more than samples with low ground kernel ash content (Figure 4A). A t-test between the high and low ground kernel ash content groups was significant (p=1.89×10), validating the importance of this trait. Ground kernel ash content is the quantity of minerals in a sample (Pojić et al., 2010), and may be representing binders or stabilizers that make pericarp more recalcitrant to base hydrolysis. Ground kernel ash content could also be an indicator of another trait that is not as easily quantified by NIR spectroscopy such as lignin content, hydroxyproline rich glycoproteins, or proline-rich proteins, each of which are known to be present in the pericarp of maize (Damasceno Junior et al., 2022) and are known to associate with mineral elements (Dar et al., 2016; Lamport & Várnai, 2013; Lao et al., 2023). Further research is needed to fully understand what biological and chemical mechanisms underlie the relationship between pericarp retention and ground kernel ash content.

**Figure 4.**
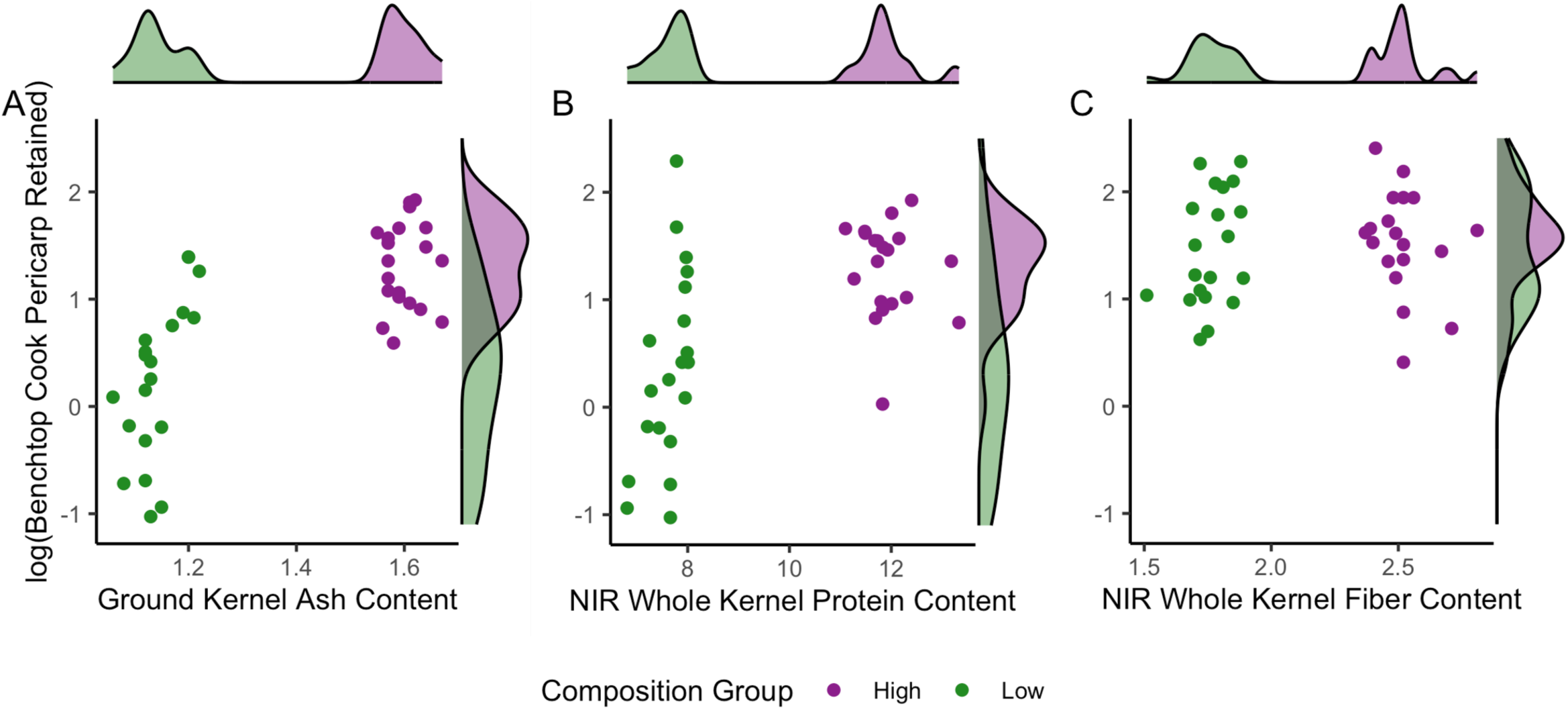
Validation of compositional relationship with pericarp retention. Twenty samples from the high and low tail of each compositional trait of interest that did not overlap to initial 60 samples was selected for validation. (A) High and low ground kernel ash content (%) samples were compared for log transformed benchtop cook test pericarp retention (mg/g). (B) High and low whole kernel protein content (%) samples were compared for log transformed benchtop cook test pericarp retention (mg/g). (C) High and low ground kernel fiber content (%) samples were compared for log transformed benchtop cook test pericarp retention. The distribution of all traits are shown on their respective axes with color associated to whether the sample was a high or low composition sample.

High whole kernel protein samples had an average of 1.81mg of pericarp per gram of kernel more than low whole kernel protein samples (Figure 4B). This difference in group mean was significant (p=1.61×10) in the independent sample set, validating that whole kernel protein content is related to nixtamalization pericarp retention. Proteins are known to be present in the cell wall of plant cells, which is the primary constituent of the pericarp (Bressani et al., 1989; Hood et al., 1991). These proteins can act as binders for the hemicellulose and cellulose chains (Damasceno Junior et al., 2022), potentially leading to increased stability during degradation processes like nixtamalization. While further research is required to identify proteins that impact pericarp retention during nixtamalization, it is worth considering proteins that interact with mineral ions as both ground kernel ash content and whole kernel protein content were significantly associated with pericarp retention in independent sample sets.

Unlike ground kernel ash content and whole kernel protein content, ground kernel fiber content was not significantly associated with pericarp retention in an independent sample set (p=0.304; Figure 4C). This result is surprising given that a large compositional component of pericarp is fiber (Damasceno Junior et al., 2022; Rocha-Villarreal et al., 2021). The term fiber is a broad term that includes cellulose, hemicellulose, and bound constituents like p-coumaric acid or ferulic acid (Chateigner-Boutin et al., 2016; Damasceno Junior et al., 2022; Machinet et al., 2011), and it is possible that if assessed individually, a relationship may be observed. In addition, the fiber content of these samples was measured on ground kernels rather than whole kernels due to equation availability. While the pericarp is thought of as a primary source of fiber in kernels, it is not the only source, and fiber content in the endosperm and germ of a kernel could be adding noise to the relationship between pericarp fiber and pericarp retention.

Further evaluation of pericarp composition is needed to fully elucidate the effects of pericarp composition in pericarp retention. Defining the role of ash content could be instrumental in breeding for grain with ideal mineral composition. Determining which proteins affect pericarp retention could give manufacturers a way to quickly test for quantity of pericarp retention if ideal protein composition is met. Lastly, determining if other aspects of pericarp composition, such as lignins or hemicellulose to cellulose ratios impact pericarp retention is vital to understanding the biology and chemistry underlying nixtamalization pericarp retention.

### 3.6 Initial pericarp quantity directly correlates with pericarp retention

From the kernel morphology traits, initial pericarp quantity and kernel circularity were further assessed to validate the potential relationship with pericarp retention. Initial pericarp quantity measures the mass of pericarp on a raw kernel and circularity measures the ratio of the kernel perimeter to the kernel area. Both traits are prohibitively time-consuming to measure across the entire panel of diverse hybrid samples. Thus, sampling the tails of the initial pericarp quantity distribution to perform segregation analysis, as was done with validating the compositional traits, was not possible. Instead, a subset of 200 spectrally diverse samples that did not have overlapping egg parents with the initial 60 sample subset was identified. Initial pericarp quantity and circularity were measured on the 200 samples, and 40 samples that spanned the range of initial pericarp quantity and kernel circularity were cooked in the benchtop cook test to quantify pericarp retention (Supplemental Figure S8; Supplemental Table S7). The Pearson’s correlation coefficient of benchtop cook test pericarp retention with initial pericarp quantity was 0.47 (p=0.0024; Figure 5A) and with kernel circularity was -0.013 (p=0.94; Figure 5B). The significant relationship with initial pericarp quantity validates the relationship between initial pericarp quantity and pericarp retention. The lack of significant relationship with kernel circularity indicates that the correlation and model importance were likely spurious, but could also be due to the fact that the range of circularity found in the initial 60 samples was broader (0.61 to 0.85) than in the validation dataset (0.68 to 0.84) and the correlation in the initial 60 samples appears to be largely driven by pericarp retention of samples with lower kernel circularity.

**Figure 5.**
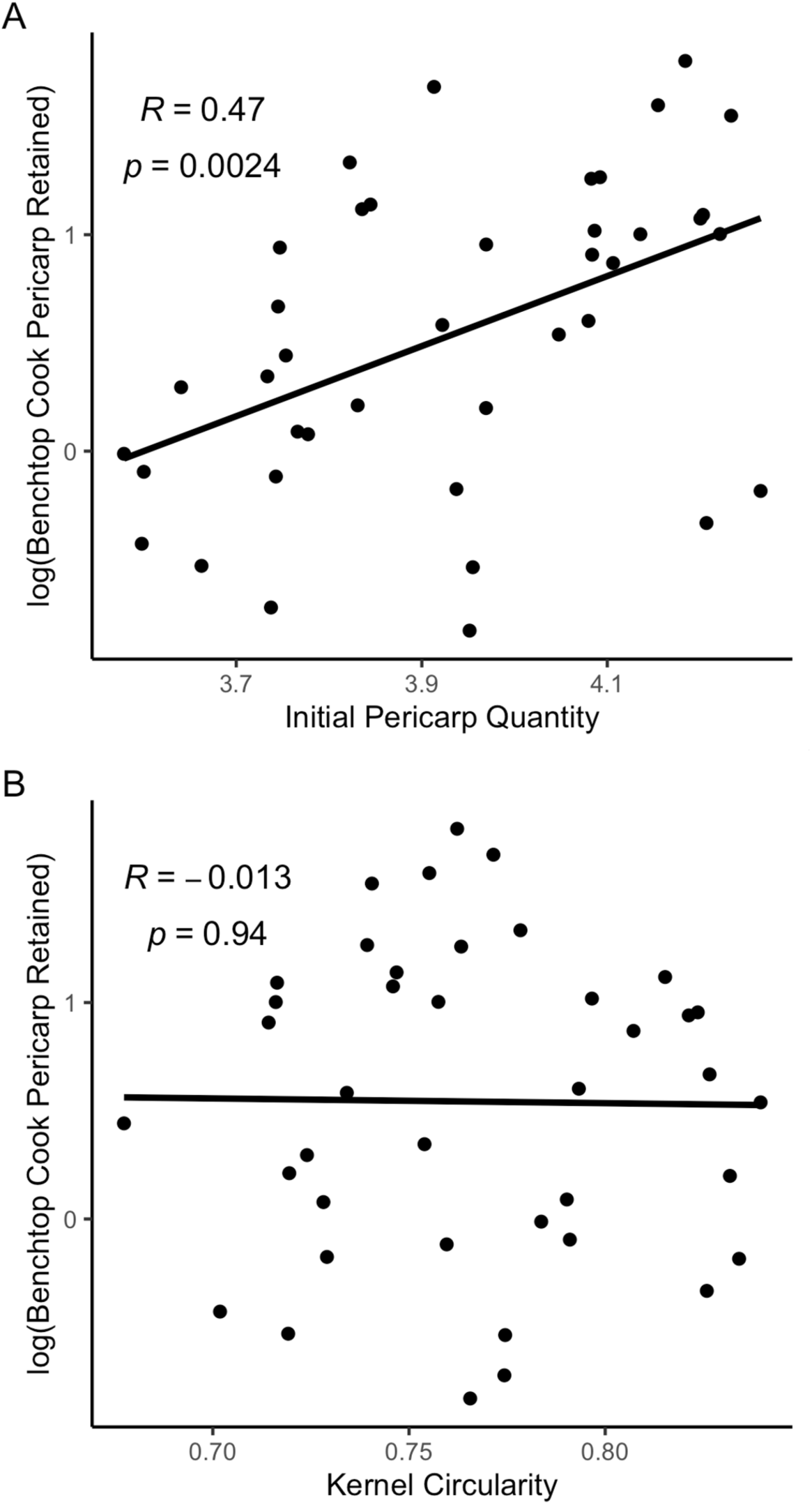
Validation of initial pericarp quantity and circularity relationship with pericarp retention. (A) Correlation of initial pericarp quantity (mg/g) and log transformed pericarp retention (mg/g). (B) Correlation of circularity and log transformed pericarp retention (mg/g). Samples chosen for validation include forty samples that evenly distributed across the range of initial pericarp quantity and circularity of 200 spectrally diverse hybrids

The importance of initial pericarp quantity is somewhat unsurprising as more substrate will inherently take longer to break down by base hydrolysis (Iribarren et al., 2002). However, there are many aspects of initial pericarp quantity that remain unknown. For example, pericarp is thinner near the crown of the kernel than the tip of the kernel (Chateigner-Boutin et al., 2016), and how the change in pericarp thickness across the kernel impacts pericarp retention is unknown. Initial pericarp quantity is also largely driven by maternal genetics and impacted by environmental growing conditions (Chateigner-Boutin et al., 2016), which could affect the cellulose/hemicellulose ratios, and even the type of hemicellulose present in the pericarp (Shilong Zhang et al., 2022). These changes could alter the density of the pericarp, which would in turn affect the mass of the initial pericarp quantity, even if the volume of pericarp does not change. More research is needed to better understand the biological basis of the effect of initial pericarp quantity on pericarp retention.

## 4 CONCLUSION

This study assessed a stain rating system for ease of pericarp removal as a proxy for quantifying the mass of pericarp retained after nixtamalization, assessed a rapid cook test methodology as a higher-throughput alternative to the benchtop cook test method for quantifying pericarp retention, and created a biological foundation for future improvement of nixtamalization pericarp retention. We showed that in a spectrally diverse subset of a diverse hybrid maize panel the stain rating method was only moderately correlated with the mass of pericarp retained, and lacked the ability to discriminate between rating groups at extreme values. It was also demonstrated that pericarp retention measured through the rapid cook test is not a valuable proxy for benchtop cook test pericarp retention, which more closely resembles the expected pericarp retention in a manufacturing facility. These findings suggest that direct measurement of pericarp retention in a representative nixtamalization method is necessary to accurately quantify pericarp retention. The relationship of many kernel composition and morphology traits were tested against benchtop cook test pericarp retention and showed a strong relationship with initial pericarp content, ground kernel ash content, and whole kernel protein content. These relationships were further validated in an independent set of hybrids. These results highlight the complexity of factors contributing to pericarp retention and provide kernel attributes to consider prior to nixtamalization. The findings of this study lay the groundwork for future research into the biology and chemistry of pericarp retention and by extension, masa quality.

## ACKNOWLEDGEMENTS

We thank Moriah Weiss and Avery Rahe for their diligent work collecting the pericarp quantity data used in this study. We also thank the Minnesota Supercomputing Institute (MSI) at the University of Minnesota for providing resources that contributed to the research results reported in this paper. This work was funded by PepsiCo, Inc. The views expressed in this manuscript are those of the authors and do not necessarily reflect the position or policy of PepsiCo, Inc. MJB was funded by the University of Minnesota Informatics Institute MnDRIVE Graduate Assistantship and the University of Minnesota Doctoral Dissertation Fellowship.

## AUTHOR CONTRIBUTIONS

CNH conceived this experiment. MJB, SPB, PJH, AMG conducted the experiments. MJB and SPB analyzed the data and visualized the data. MJB and CNH wrote the original draft. All co-authors edited and approved the final manuscript.

## SUPPORTING INFORMATION

The data that supports the findings of this study are available in the supplementary material of this article.

## Notes

### Competing Interest Statement

The authors have declared no competing interest.

