## Supplemental Figures for "Maize kernel composition and morphology influences pericarp retention during nixtamalization"

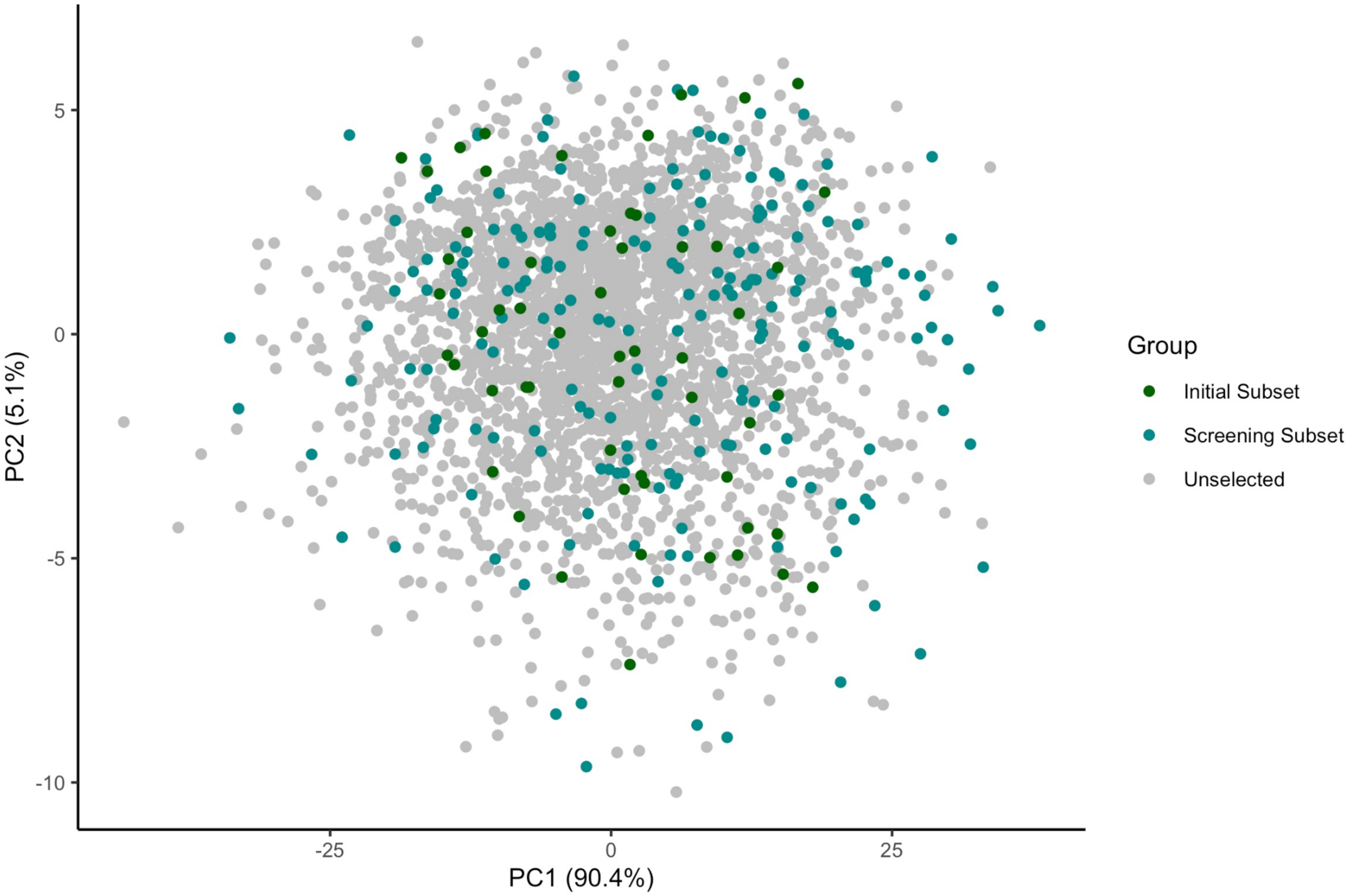


**Figure S1. Spectral diversity of Wisconsin Diversity Panel hybrids.** Spectral diversity was determined through principal component analysis of the whole kernel spectra. Samples are colored by their use in this study. Initial subset samples are the 60 spectrally diverse samples used to compare benchtop cook test pericarp retention to the rapid cook test method, stain rating methods, kernel composition, and kernel morphology. Screening subset samples were 200 samples used to evaluate initial pericarp quantity on a diverse set of germplasm for trait validation.

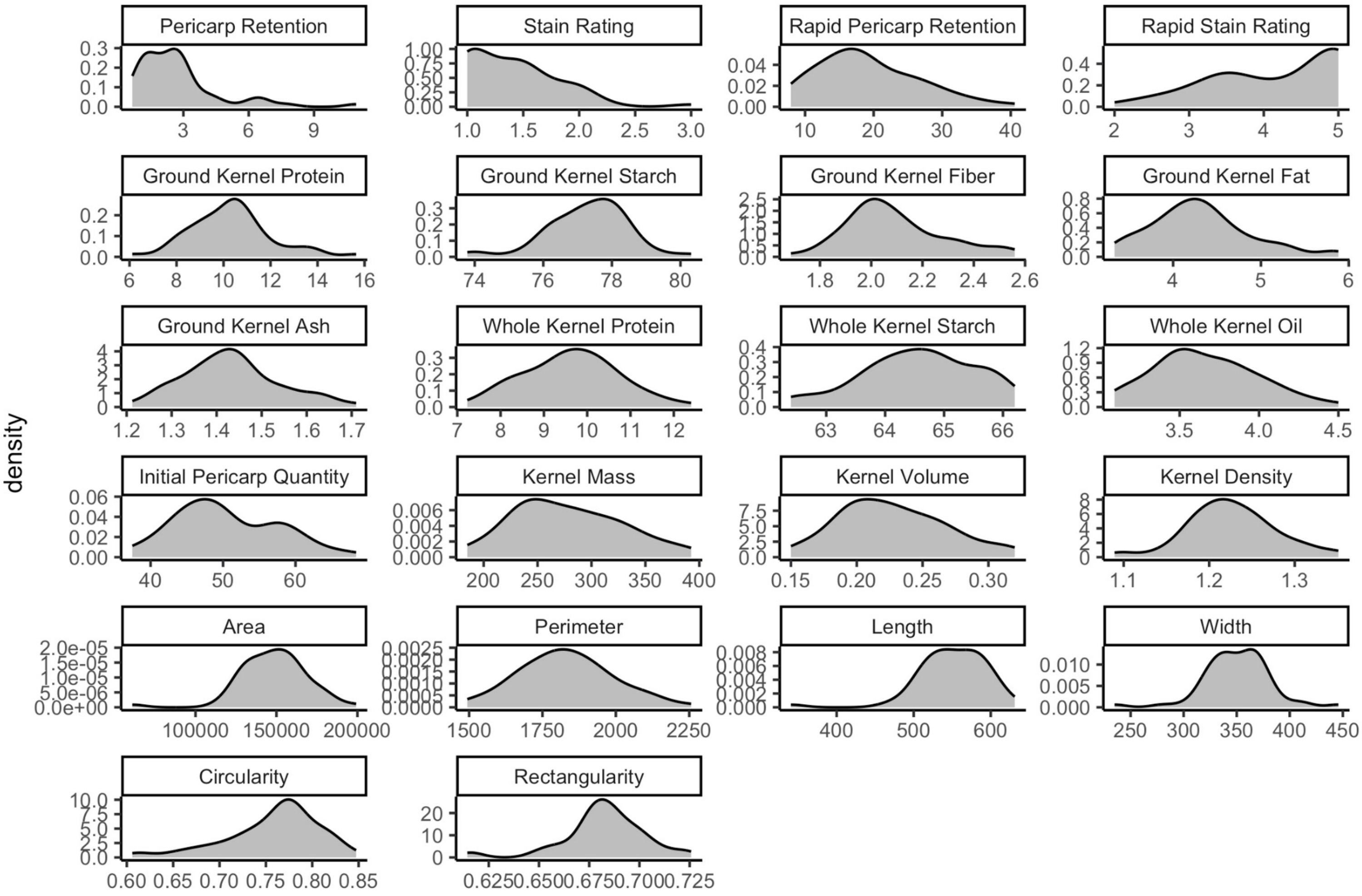


**Figure S2. Distribution of traits collected for 60 spectrally diverse hybrids.** Traits were collected through benchtop nixtamalization cook tests (Pericarp Retention (mg/g), Image Rating), rapid nixtamalization cook tests (Rapid Pericarp Retention, Rapid Image Rating), Perten NIR spectroscopy (Ground Kernel Protein (%), Ground Kernel Starch (%), Ground Kernel Fiber (%), Ground Kernel Fat (%), Ground Kernel Ash (%)), FOSS NIR spectroscopy (Whole Kernel Protein (%), Whole Kernel Starch (%), Whole Kernel Oil (%)), manual measurement (Initial Pericarp Quantity (mg/g), Kernel Mass (g), Kernel Volume (ml), Kernel Density (g/ml)), and image analysis (Area (px), Perimeter (px), Length (px), Width (px), Circularity, Rectangularity).

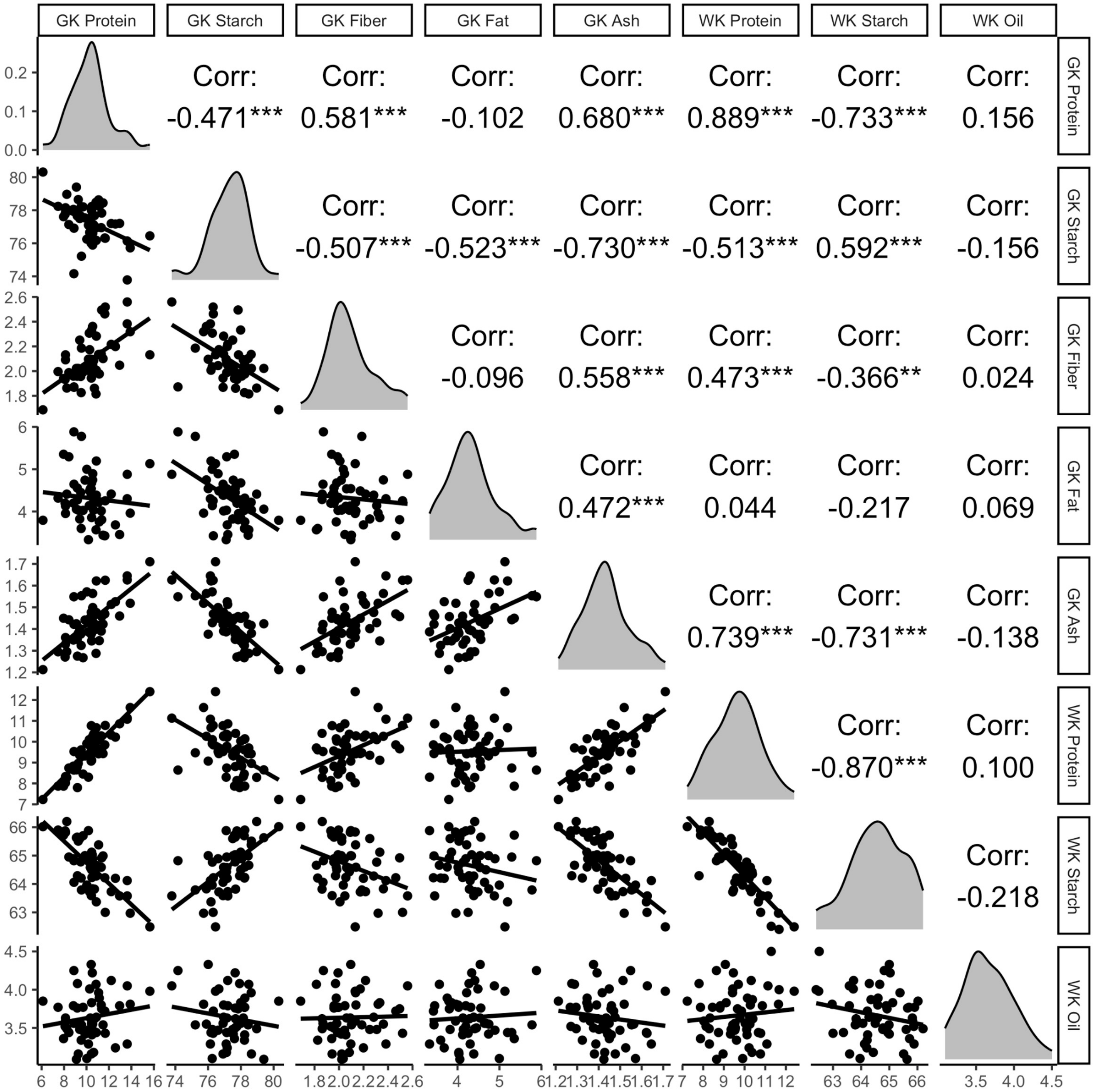


**Figure S3. Distribution and correlation and compositional traits collected from Perten and FOSS NIR spectrometers.** Pearson's correlation coefficient was determined between all pairwise correlations of compositional traits collected from ground kernel (GK) samples on a Perten DA7250 and whole kernel (WK) samples on a FOSS Infratec Nova.

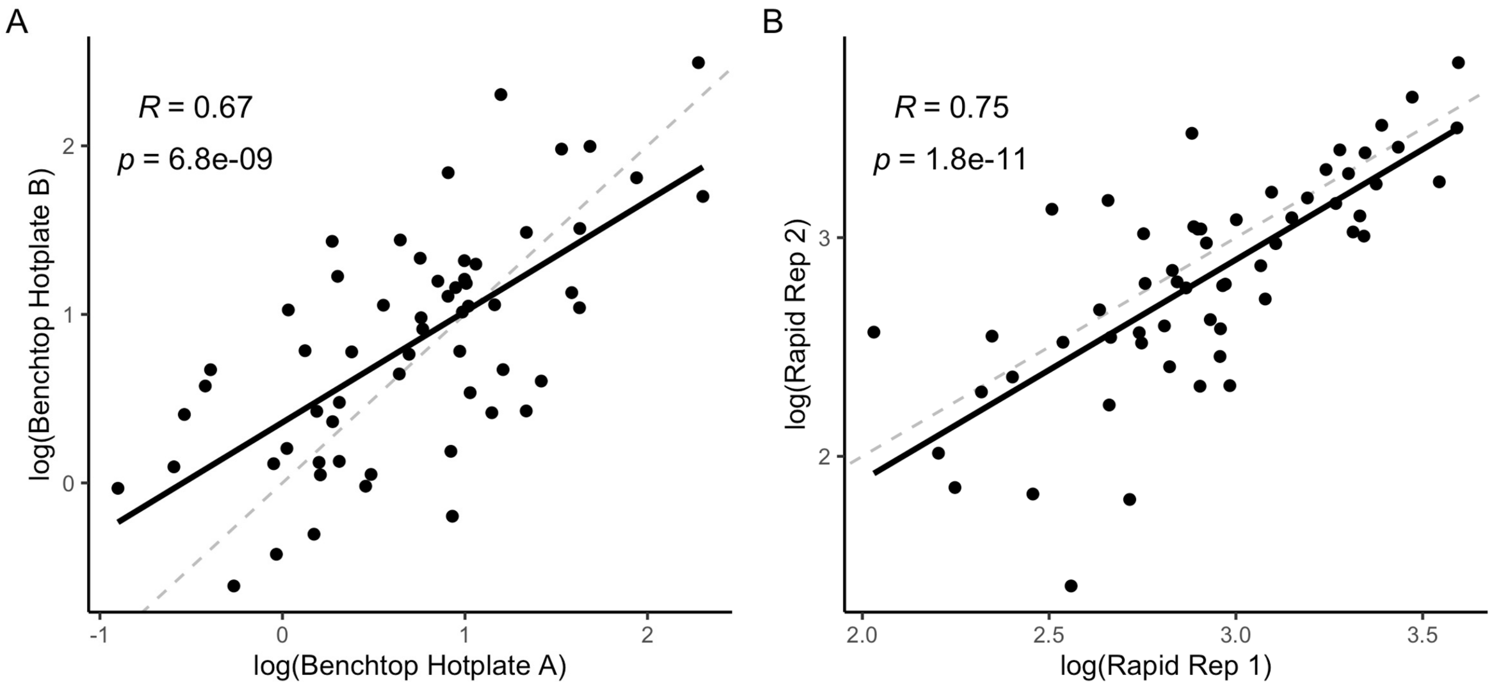


**Figure S4. Correlation of biological replicates in the benchtop and rapid cook test methods.** (A) Correlation of log transformed pericarp retention (mg/g) from replicates cooked across hotplates in the benchtop cook test. (B) Correlation of log transformed pericarp retention (mg/g) from replicates cook in the rapid cook test.

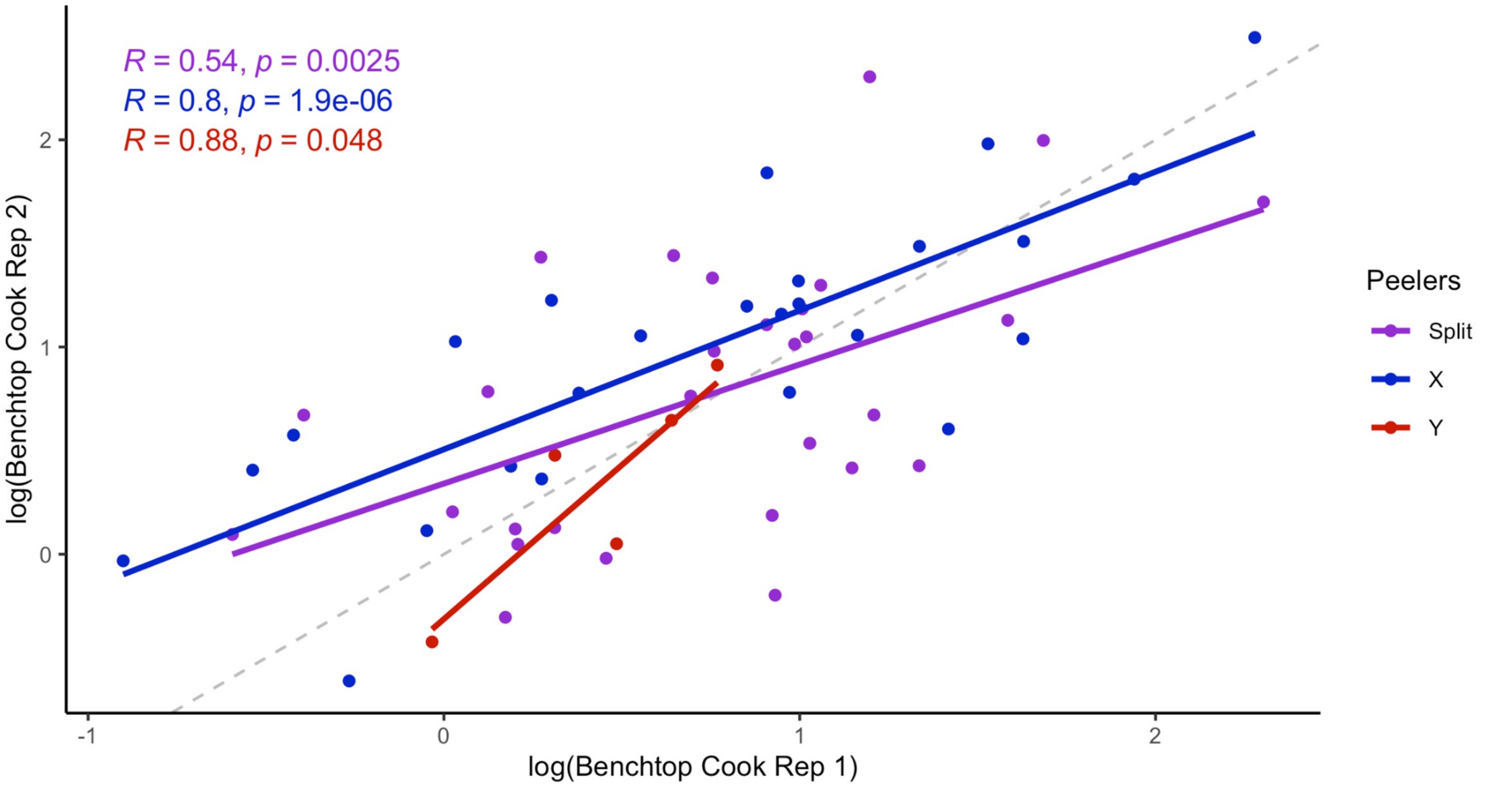


**Figure S5. Comparison of peeler consistency in the benchtop cook test.** Log transformed pericarp retention from the benchtop cook test method colored by the peeler of both replicates. Samples of peeler A or B were peeled by a single individual and samples of peeler Split were samples where one replicate was peeled by peeler A and the other replicated peeled by peeler B.

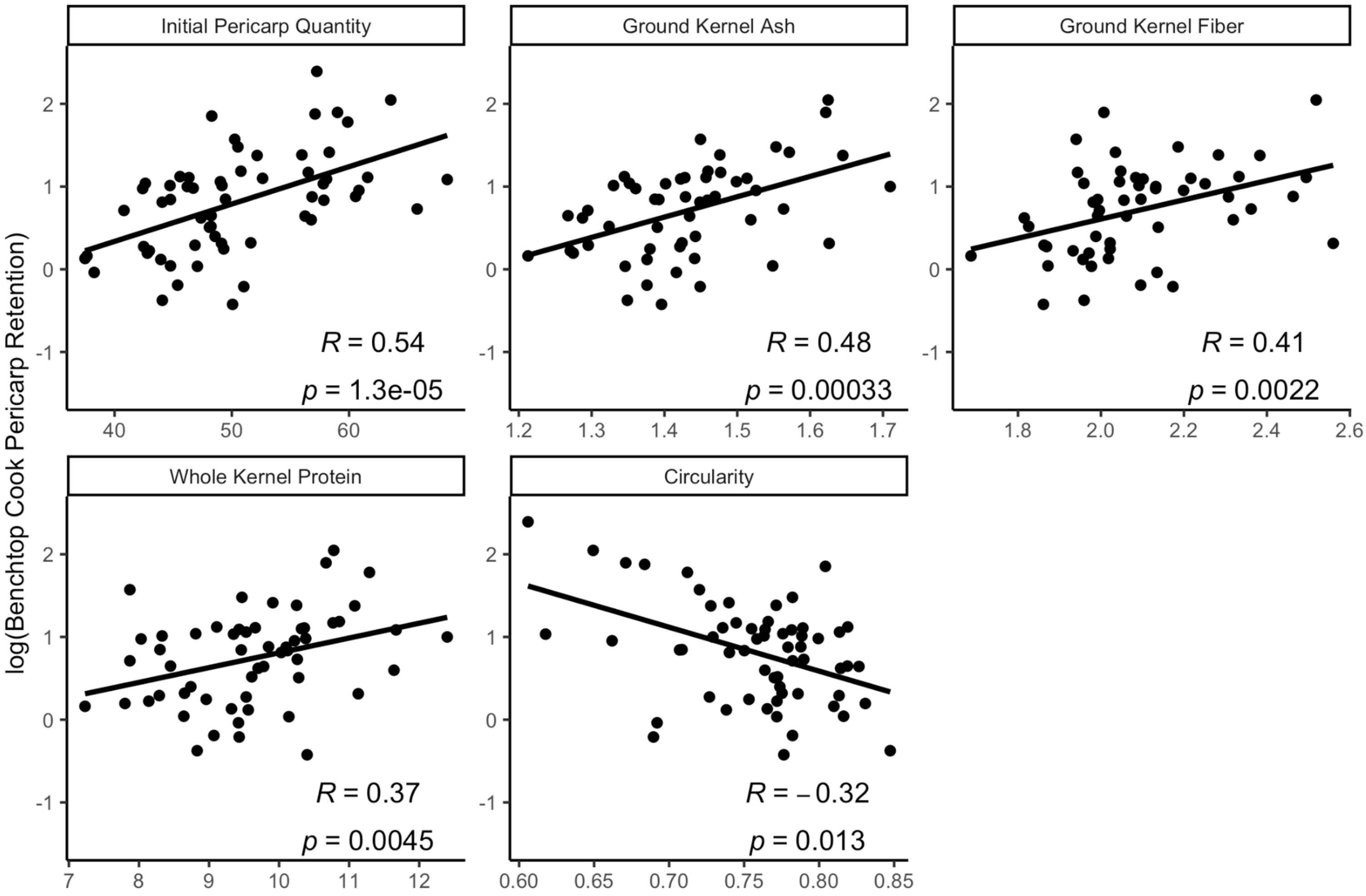


**Figure S6. Correlation plots of the most associated traits with pericarp retention from the initial 60 sample set.** The correlation of initial pericarp quantity (mg/g), ground kernel ash content (%), circularity, ground kernel fiber content (%), and whole kernel protein content (%) with log transformed benchtop cook test pericarp retention (mg/g). Correlation is calculated with Pearson's correlation coefficient.

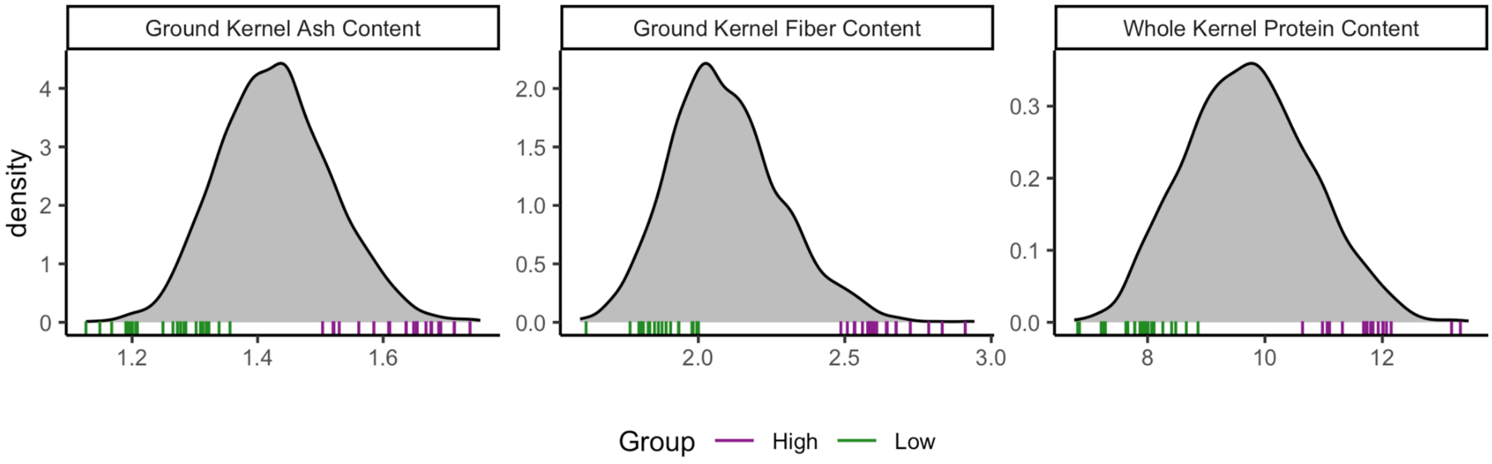


**Figure S7. Distribution of compositional traits from remaining Wisconsin Diversity Panel hybrids for validation.** The distribution of all Wisconsin Diversity Panel hybrids not in the initial 60 sample set was used to find 20 high and 20 low samples for ground kernel ash content (%), ground kernel fiber content (%), and whole kernel protein content (%) shown in tick marks colored by group association below the distribution.

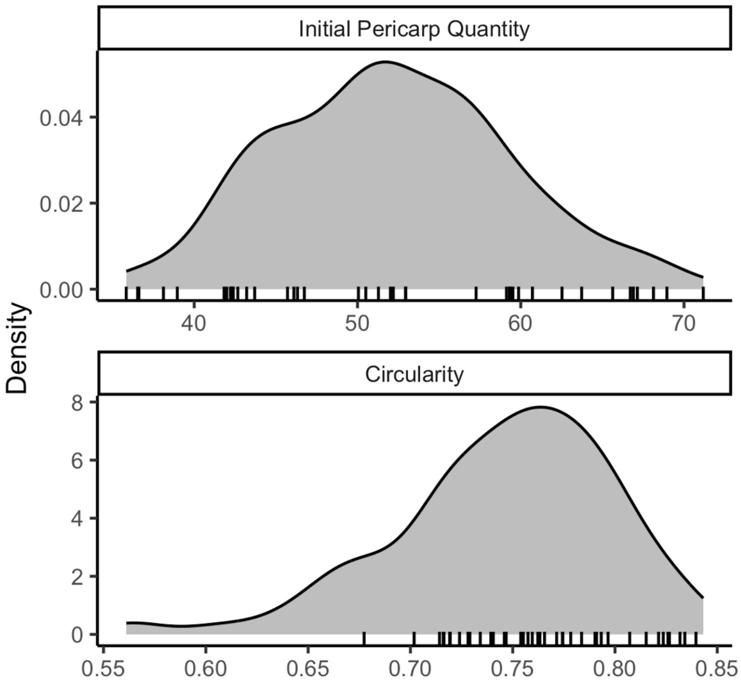


**Figure S8. Distribution of morphological traits from remaining Wisconsin Diversity Panel hybrids for validation.** The distribution of all Wisconsin Diversity Panel hybrids in the 200 screening sample set used to find 40 samples evenly spread throughout the distribution of initial pericarp quantity (mg/g) and circularity, shown in black tick marks below the distribution, which did not have overlapping egg parents with the initial set of 60 samples or within this population.
